# Shear stress-induced pathological changes in endothelial cells occur through Piezo1 activation of TRPV4

**DOI:** 10.1101/2020.07.01.182212

**Authors:** Sandip M. Swain, Rodger A. Liddle

## Abstract

Although the ion channels Piezo1 and TRPV4 have been implicated in high venous pressure- and fluid shear stress-induced vascular hyperpermeability, they have been described as working independently. Moreover, the mechanism by which Piezo1 and TRPV4 channels in endothelial cells execute the same function is poorly understood. Here we demonstrate that Piezo1 regulates TRPV4 channel activation in endothelial cells and that Piezo1-mediated TRPV4 channel opening is a function of the strength and duration of fluid shear stress. Application of the Piezo1 antagonist, GsMTx4, completely blocked the elevation in intracellular calcium ([Ca^2+^]_i_) induced by both fluid shear stress and the Piezo1 agonist, Yoda1. High and prolonged shear stress caused sustained [Ca^2+^]_i_ elevation which required TRPV4 opening and was responsible for fluid shear stress- and Piezo1-mediated disruption of adherens junctions and actin remodeling. We found that Piezo1’s effects were mediated by phospholipase A2 activation. Blockade of TRPV4 channels with the selective TRPV4 blocker, HC067047, prevented the loss of endothelial cell integrity and actin disruption induced by Yoda1 or shear stress and prevented Piezo1-induced monocyte adhesion to endothelial cell monolayers. These findings demonstrate that Piezo1 activation by fluid shear stress initiates a calcium signal that causes TRPV4 opening which in turn is responsible for the sustained phase calcium elevation that triggers pathological events in endothelial cells.

## Introduction

Endothelial cells are under constant mechanical force due to blood pressure and flow in both the arterial and venous systems. Blood flowing across endothelial cells within blood vessels generates shear stress. Under normal physiological conditions, a dynamic balance between mechanical shear stress and biological responses maintains endothelial integrity (1-5). Shear stress induces the release of endothelial vasodilatory factors such as nitric oxide, prostacyclin, and cytochrome *P-450* metabolites of arachidonic acid that are required for underlying smooth muscle relaxation (6-8). Endothelial Ca2+-mediated mechanosensing is required for normal flow-mediated dilation (4,5). Perturbation of endothelial shear stress that occurs with hypertension or excessive flow leads to vascular remodeling through the disruption of cytoskeletal proteins and vascular dysfunction involving the loss of endothelial integrity, increased endothelial cell stiffness, altered vasorelaxation properties, and leukocyte adhesion (2,3,9-13).

Clinically, high venous pressure is a major cause of pulmonary edema and mortality in patients with congestive heart failure (10,11). Elevated vascular pressure can lead to endothelial barrier disruption and hyperpermeability due to loss of adherens junctions (AJs) between endothelial cells (3,11). It has recently been demonstrated that the mechanosensitive ion channel Piezo1 mediates pressure-induced disruption of AJs and endothelial barrier breakdown in pulmonary vessels (11,14). Piezo1 is activated by cell membrane tension caused by high pressure, shear stress and membrane stretching which allow the influx of cations, mainly Ca2+, and triggers downstream calcium signaling (15-18). These processes are important for maturation of the vasculature as deletion of Piezo1 impaired vascular development in mice and also blocked sprouting angiogenesis in response to shear stress (16). However, endothelial Piezo1 mediates pathological responses to pressue and is involved in atherosclerosis progression and inflammatory signaling (19).

Like Piezo1, the endothelial cell-expressed, calcium-permeable transient receptor potential vanilloid subfamily 4 (TRPV4) channel is expressed in various tissues and cells that are also pressure-sensitive (e.g., vascular endothelium, urinary bladder, and airway epithelium) (9,20,21). TRPV4 has been linked to several physiological functions including epithelial ciliary activity, regulation of blood flow, and shear-induced vasodilation and angiogenesis, and pathological processes that involve endothelial dysfunction and actin disruption (4,22-24). Blockade of TRPV4 channels protects against pulmonary edema and hyperpermeability induced by high pressure (10). TRPV4 channels are believed to be activated by shear stress, membrane stretching and hypotonic cell swelling although they do not have true mechanoreceptor properties (4,7,25). Therefore, how TRPV4 in endothelial cells senses physical force is unknown.

It was demonstrated previously that blood flow-mediated shear stress activates phospholipase A2 (PLA2) generating 5′,6′-epoxyeicosatrienoic acid (5′,6′-EET) from arachidonic acid (6,26). Importantly, 5′,6′-EET has the ability to activate TRPV4. However, how shear stress activates PLA2 in endothelial cells is unknown.

We recently observed that stimulation of Piezo1 in pancreatic acinar cells is responsible for pressure-induced pancreatitis (27) and is linked to TRPV4 (28). Therefore, we postulated that Piezo1 signaling coupled to TRPV4 activation may also account for the effects of shear stress on endothelial cells. Here we demonstrate that high shear stress activates Piezo1 and causes an initial increase in [Ca^2+^]_i_ that triggers activation of TRPV4. Activation of TRPV4 causes a sustained [Ca^2+^]_i_ elevation leading to loss of endothelial cell contacts, actin disruption, and endothelial cell monocyte adhesion.

## Results

### Fluid shear stress induced [Ca^2 +^]_i_ overload is force and time dependent

Shear stress is a physiological activator of mechanical ion channels in endothelial cells. In endothelial cells, shear stress regulates levels of intracellular calcium ([Ca^2+^]_i_) and downstream calcium signaling (23,29-31). We evaluated the magnitude and duration of shear stress forces that affected [Ca^2+^]_i_ in human umbilical vein endothelial cells (HUVECs). Shear stresses of 4 and 12 dyne/cm^2^ for 1 min increased peak [Ca^2+^]_i_ (Fig. 1A, and B). Applying a force of 12 dyne/cm2 for 1 min caused a sustained elevation in [Ca^2+^]_i_ (intensity calculated at 8 min after initiation of force) but the same force applied for a shorter time (12 dyne/cm2 for 5 seconds) or lower shear stress (4 dyne/cm2) for 1 min elicited only a transient [Ca^2+^]_i_ rise and did not produce prolonged elevation in [Ca^2+^]_i_ (Fig. 1A, B and C).

**Figure 1.**
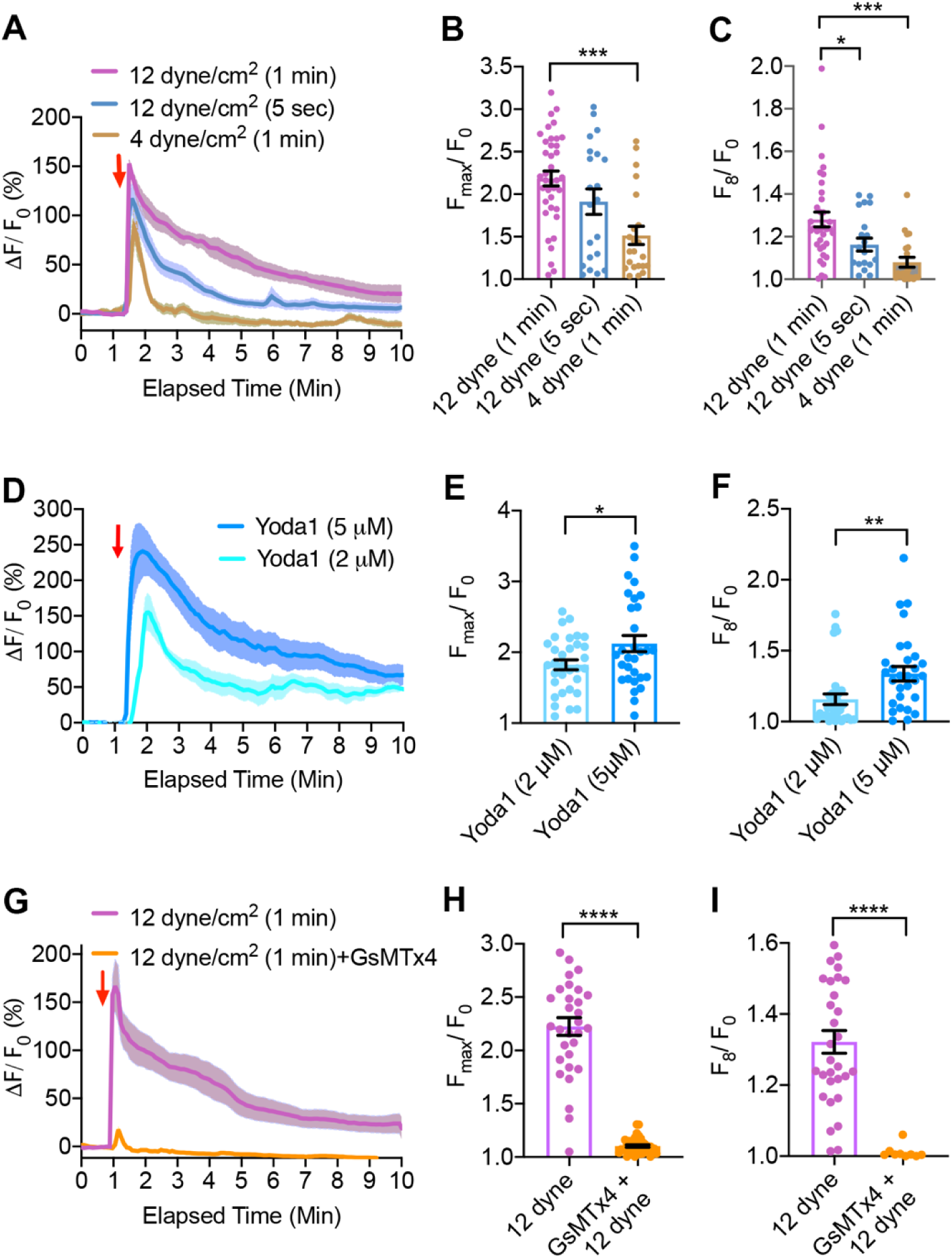
Fluid shear stress and Yoda1 induced [Ca^2+^]i elevation in HUVECs. (A) The relative fluorescence intensity (ΔF/F0) of calcium 6-QF–loaded HUVECs are shown in response to applied shear stress 12 dyne/cm2 or 4 dyne/cm2 for 1 min or 5 sec. (B) The graphs show the average maximum peak [Ca2+]i intensity (Fmax/F0), and (C) the average peak intensity calculated at 8 min after initiation of force from 21–35 cells, n = 3–4 independent experiments. (D, E and F) Yoda1-induced [Ca2+]i elevation in HUVECs. (D) The relative fluorescence intensity (ΔF/F0) of calcium dye over time with Yoda1 (2 μM) or Yoda1 (5 μM). (E) The graphs show the average maximum peak [Ca2+]i intensity and (F) the average [Ca2+]i intensity peak calculated at 8 min after the initiation of force from 30 to 32 cells. (G, H and I) The effects of GsMTx4 on fluid shear stress-induced [Ca2+]i elevation. GsMTx4 was applied 2 min before shear stress. (G) The relative fluorescence intensity (ΔF/F0) of calcium dye overtime with shear stress 12 dyne/ cm2 for 1 min with or without GsMTx4 (10 μM). (H) The average maximum peak [Ca2+]i intensity, and (I) the average [Ca2+]i intensity peak calculated at 8 min after initiation of force from 31 to 40 cells, n = 3 experiments. Statistical analyses were performed using two-tailed Student’s t test *P ≤ 0.05; **P ≤ 0.01; ***P ≤ 0.001; ****P ≤ 0.0001, Data are shown as mean±SEM.

Endothelial cells express the mechanically sensitive, calcium-permeable ion channel, Piezo1(14,32). To evaluate the role of Piezo1 channels in sensing shear force, we utilized the Piezo1 agonist, Yoda1. Yoda1 (2 μM and 5 μM), in a dose-dependent manner, increased peak [Ca^2+^]_i_, and caused sustained [Ca^2+^]_i_ elevations (fluorescence intensity calculated at 8 min after Yoda1 application) (Fig. 1D, E and F). Thus, the level of [Ca^2+^]_i_ produced by Piezo1 activation in HUVECs was dependent upon the Yoda1 concentration (Fig. 1D, E and F). To determine if Piezo1 was responsible for shear stress-induced calcium influx, we used the mechanoreceptor blocker, GsMTx4 (33). GsMTx4 (10 μM) completely inhibited the shear stress (12 dyne/cm^2^)-induced elevation in [Ca^2+^]_i_ (Fig 1G, H and I).

### TRPV4 is responsible for the Piezo1-induced sustained [Ca^2+^]_i_ elevation

Although Piezo1 directly senses mechanical force (18), its fast inactivation kinetics and low single channel conductance (15), render it unlikely to be directly responsible for the sustained elevation in [Ca^2+^]_i_ seen with higher shear forces. It is notable that endothelial cells also express another calcium-permeable ion channel - TRPV4 (4,30). Even though cells expressing TRPV4 exhibit mechanosensitivity, direct channel activation by mechanical force has not been demonstrated (9). Therefore, the ability of shear stress to activate TRPV4 channels seems to be indirect. To evaluate the contribution of TRPV4 to sense shear force, we tested 1 μM HC067047 (HC067), a concenration that has been reported to completely block TRPV4 channel activity (21). HC067 slightly reduced the initial rise in [Ca^2+^]_i_ produced by shear force applied at 12 dyne/cm^2^ for 1 min but completely blocked the sustained calcium elevation (Fig. 2A, B and C). This observation demonstrates that activation of the TRPV4 channel is responsible for the secondary, sustained elevation in [Ca^2+^]_i_ produced by shear force. Addtionally, HC067(1 μM) completely blocked the Yoda1 induced sustained [Ca^2+^]_i_ rise without affecting initial rise in [Ca^2+^]_i_ confirming that Yoda1 and shear stress induced secondary [Ca^2+^]_i_ rise phase is due to activation of TRPV4 and initial transient [Ca^2+^]_i_ rise through activation of Piezo1 (Fig. 2, D, E and F).To determine if selective activation of TRPV4 channels can reproduce the sustained [Ca^2+^]_i_ elevation, we tested the effects of theTRPV4 agonist, GSK1016790A (GSK101) and endogenous TRPV4 agonist 5′,6′-epoxyeicosatrienoic acid (5,6 -EET). Both GSK101 (50 nM) and 5,6 -EET (5 μM) produced sustained [Ca^2+^]_i_ elevations in HUVECs (Fig. 2G, H and I).

**Figure 2.**
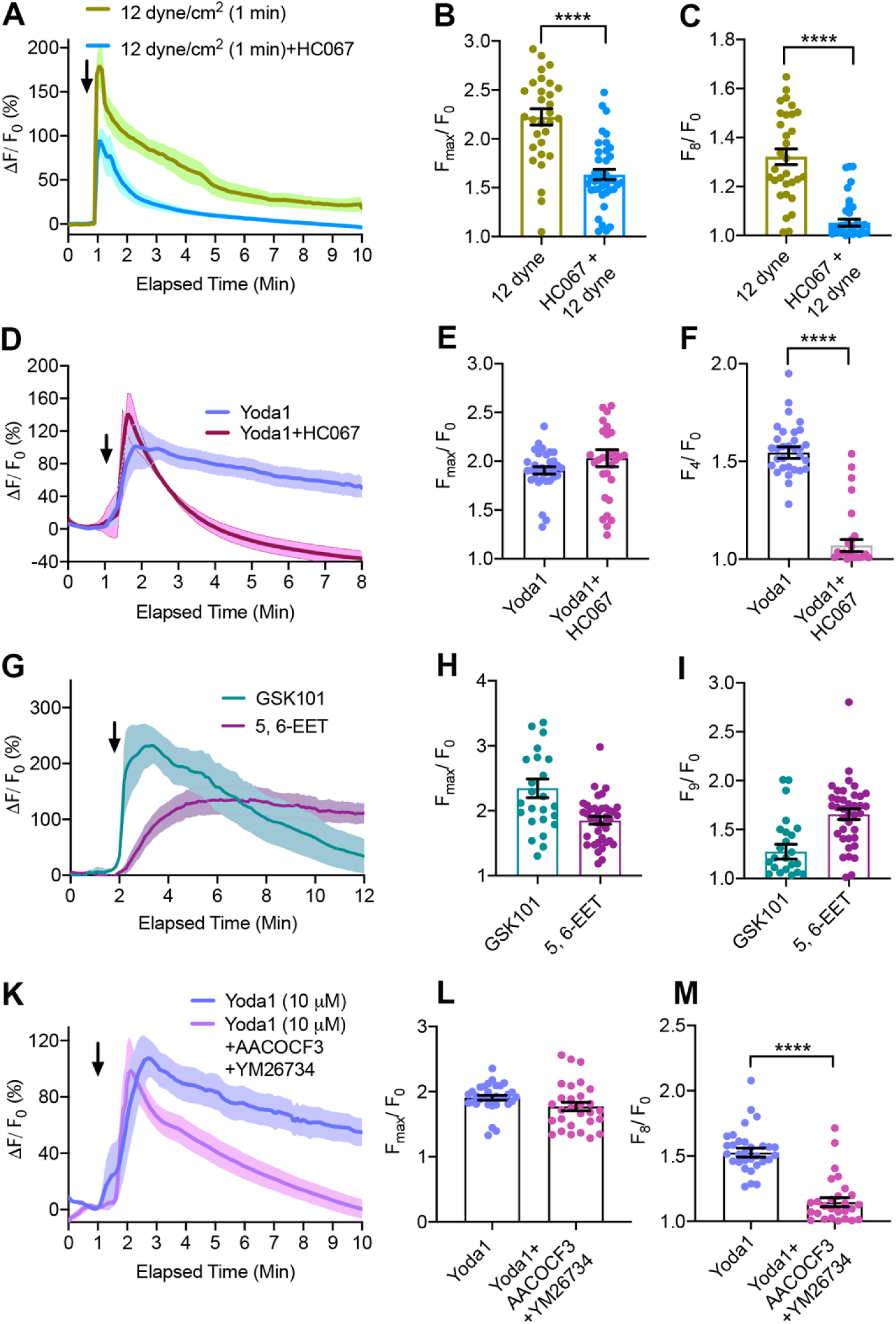
HC067 and PLA2 blockers inhibited the Piezo1-induced sustained [Ca^2+^]i elevation in HUVECs. (A, B and C) The effects of HC067 on fluid shear stress-induced [Ca2+]i elevation. HC067 was applied 2 min before shear stress. (A) The relative fluorescence intensity (ΔF/F0) of calcium dye overtime with shear stress 12 dyne/ cm2 for 1 min with or without HC067 (1 μM). (B) The average maximum peak [Ca2+]i intensity and (C) the average peak [Ca2+]i intensity calculated at 8 min after the force from 31 to 40 cells, n = 3 experiments. (D, E, and F) The effect of HC067 (1 μM) on Yoda (5 μM) induced [Ca2+]i elevation. (D) Relative fluorescence intensity (ΔF/F0) of calcium dye over time. (E and F) Average maximum peak intensity and average peak intensity at 4 min from 29-32 cells from three experiments. (G, H and I) GSK101 (50 nM) and 5′,6′-EET (5 μM) increased [Ca2+]i. (G) The relative fluorescence intensity of calcium dye, (H) average maximum peak [Ca2+]i intensity, and (I) the average [Ca2+]i intensity peak calculated at 9 min after initiation of force from 25 to 38 cells. (K, L and M) The effects of PLA2 blockers, AACOCF3 (30 μM) and YM26764 (10 μM) on Yoda1 (10 μM) induced [Ca2+]i rise. (K) The relative fluorescence intensity of calcium dye overtime with Yoda1 with or without AACOCF3 (30 μM) + YM26764(10 μM). AACOCF3 (20 μM) + YM26764(10 μM) were applied 8 min before Yoda1 application. (L and M) The average maximum peak intensity and average peak intensity at 8 min from 30-31 cells from three experiments. Black arrows show the time stimuli were applied. Statistical analyses were performed using two-tailed Student’s t test ****P ≤ 0.0001, Data are shown as mean ± SEM.

Prolonged exposure of HUVECs to the selective Piezo1 agonist Yoda1 induced a prolonged increase in [Ca^2+^]_i_ (Fig. 2K, L, M) raising the possibility that Piezo1 activation is coupled to TRPV4 channel opening. Intracellular signals including PLA2 have been shown to activate TRPV4 (6) and we and others have previously reported that shear stress can increase PLA2 activity (28,34). Therefore, to determine the intracellular process that is responsible for the effects of Piezo1 on TRPV4 channel opening, we applied PLA2 blockers and measured the effects of Yoda1 [Ca^2+^]_i_ in HUVECs. Treatment with the cytoplasmic PLA2 blocker AACOCF3 (30 μM) (35) and secretory PLA2 blocker YM26734 (10 μM) (36) significantly inhibited the Yoda1-induced sustained elevation in [Ca^2+^]_i_ (Fig. 2K, L and M), indicating that Piezo1 regulates PLA2 activity, which is responsible for the subsequent activation of TRPV4.

### Piezo1-induced adherens junction (AJ) disruption is the consequence of TRPV4 activation

AJs consist of p120-catenin, β-catenin, and α-catenin and the transmembrane adhesive protein VE-cadherin (11,37). Activation of Piezo1 disrupts vascular AJs and increases vascular permeability (11). To determine if this disruption occurs through Piezo1 on vascular endothelial cells, we treated monolayer HUVEC cells with Yoda1 (2 μM) for 30 min and observed a reduction in VE-cadherin expression at AJs (Fig. 3A). The overall apparent width of VE-cadherin at AJs was decreased significantly with Yoda1 (2 μM) (Fig. 3C). A higher concentration of Yoda1 (5 μM) decreased the accumulation of VE-cadherin and disrupted HUVEC integrity (Fig. 3A and B). We proposed that these changes were due to Piezo1-triggered TRPV4 activation causing secondary calcium overload. To determine if the effect of Yoda1 on VE-cadherins was due to activation of TRPV4, we evaluated the action of HC067 (1 μM). Treatment with HC067 stabilized the VE-cadherin at AJs and also protected the integrity of HUVECs in monolayer culture (Fig. 3A and B). These findings indicate that Yoda1-induced loss of AJs is through the combined actions of Piezo1 and TRPV4. We observed that a high concentration of Yoda1 (10 μM) caused cell retraction and produced paracellular gaps in monolayer cultures (Fig. 3F and Movie 1). Paracellular gaps were the consequence of the reduction in the cell surface area. Cellular disruption due to Yoda1 was absent when the TRPV4 blocker HC067 was added to the media (Fig. 1F and Movie 1).

**Figure 3.**
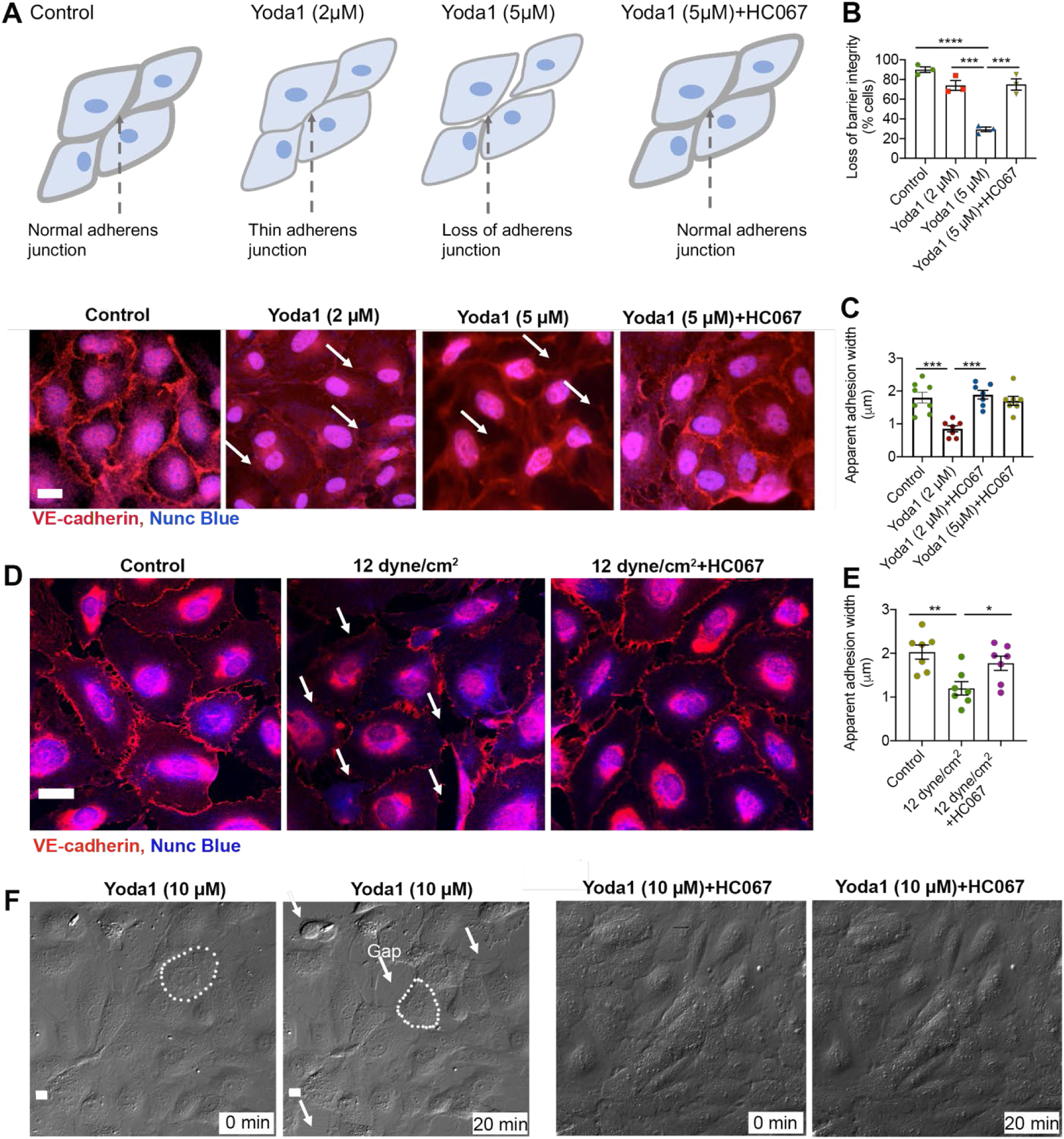
HC067 prevented Piezo1-mediated loss of AJs in HUVEC endothelial cells. (A, B, C, and D) Accumulation of the AJs protein, VE-cadherin, at junctions of HUVECs monolayer is shown by red immunostaining along the cell membrane of endothelial cells. (A) The images and cartoons show that application of Yoda1 for 30 min caused thinning (2 μM) or loss (5 μM) of VE-cadherin in endothelial cells. HC067 (1 μM) prevented Yoda1-induced loss of VE-cadherin. (B) The number of cells with loss of cellular integrity was calculated from three experiments. (C) The width of VE-cadherin (red) staining was quantified from data shown in A. (D) Effects of laminar fluid shear stress (12 dyne/cm2 for 10 min) with or without HC067 on VE-cadherin (red) accumulation at junctions of HUVECs monolayers are shown. (E) Quantification of the apparent width of VE-cadherin (red) were calculated from data shown in D. (F) Live cell DIC images of HUVECs in monolayer culture before or 20 min after Yoda1 (10 μM) or Yoda1 + HC067 (5 μM) show the loss of cellular junctions overtime, retraction of cells (a cell boundary marked in white dotted line before and after Yoda1), and formation of intracellular gaps (marked in white arrows). Individual data points are shown as mean ± SEM; Statistical analyses were performed using two tailed Student’s t test; multiple groups were analyzed by 1-way ANOVA with Tukey’s multiple comparisons. *P ≤ 0.05; ** P ≤ 0.01, *** P ≤ 0.001; ****P ≤0.0001. Scale bar: 10 μm.

Subjecting HUVEC monolayers to 12 dyne/cm2 for 10 min reduced VE-cadherin at AJs and decreased the apparent width of individual cells. Knowing that through Piezo1, shear stress triggers TRPV4 activation which is responsible for producing high levels of [Ca^2+^]_i_, we postulated that blocking TRPV4 would protect against the loss of VE-cadherin caused by Yoda1. Thus, if TRPV4 was responsible for the sustained [Ca^2+^]_i_ elevation that causes the loss of VE-cadherin, then blocking TRPV4 would be expected to protect against disruption of AJs. We observed that HC067 (1 μM) blocked shear stress-mediated loss of VE-cadherin at AJs (Fig. 3D) and preserved the apparent width of VE-cadherin (Fig. 3E).

### Inhibition of TRPV4 channels protected against shear stress-induced actin disorganization

Cytoskeleton remodeling in response to the physiological levels of shear stress is necessary for endothelial cell migration and maintaining vascular integrity (1,38). However, shear stress generated under conditions of high vascular pressure or turbulence has adverse effects on the cytoskeleton which can lead to increased endothelial cell stiffness, vascular permeability, and transmigration of leukocytes (1,11,39,40). We analyzed F-actin fiber distribution using F-actin plot profiling in which the intensity and orientation of F-actin fibers are measured (Fig. 4 A, B and C). We observed that high shear stress (25 dyne/cm2 for 10 min) disrupted the orientation of F-actin in HUVECs in contrast to lower shear force (4 dyne/cm2 for 10 min) (Fig. 4 A, B and C). High shear stress triggered the formation of F-actin bundles indicative of fiber polymerization (Fig. 4 A, B and C). The F-actin intensity frequency distribution predicted that high shear stress caused F-actin fibers to translocate from the peri-nuclear region of the cell to the periphery where a high-density of uneven clusters appeared (Fig. 4 A, B and C). The uniform thickness of F-actin fibers generally observed under low shear stress conditions was lost and extensive polymerization of thick F-actin bundles appeared at the periphery of cells subjected to high shear stress (Fig. 4). These effects of high shear forces were prevented in cells treated with GsMTx4 confirming that high shear stress-mediated F-actin disorientation required Piezo1 channel activation (Fig. 4).

**Figure 4.**
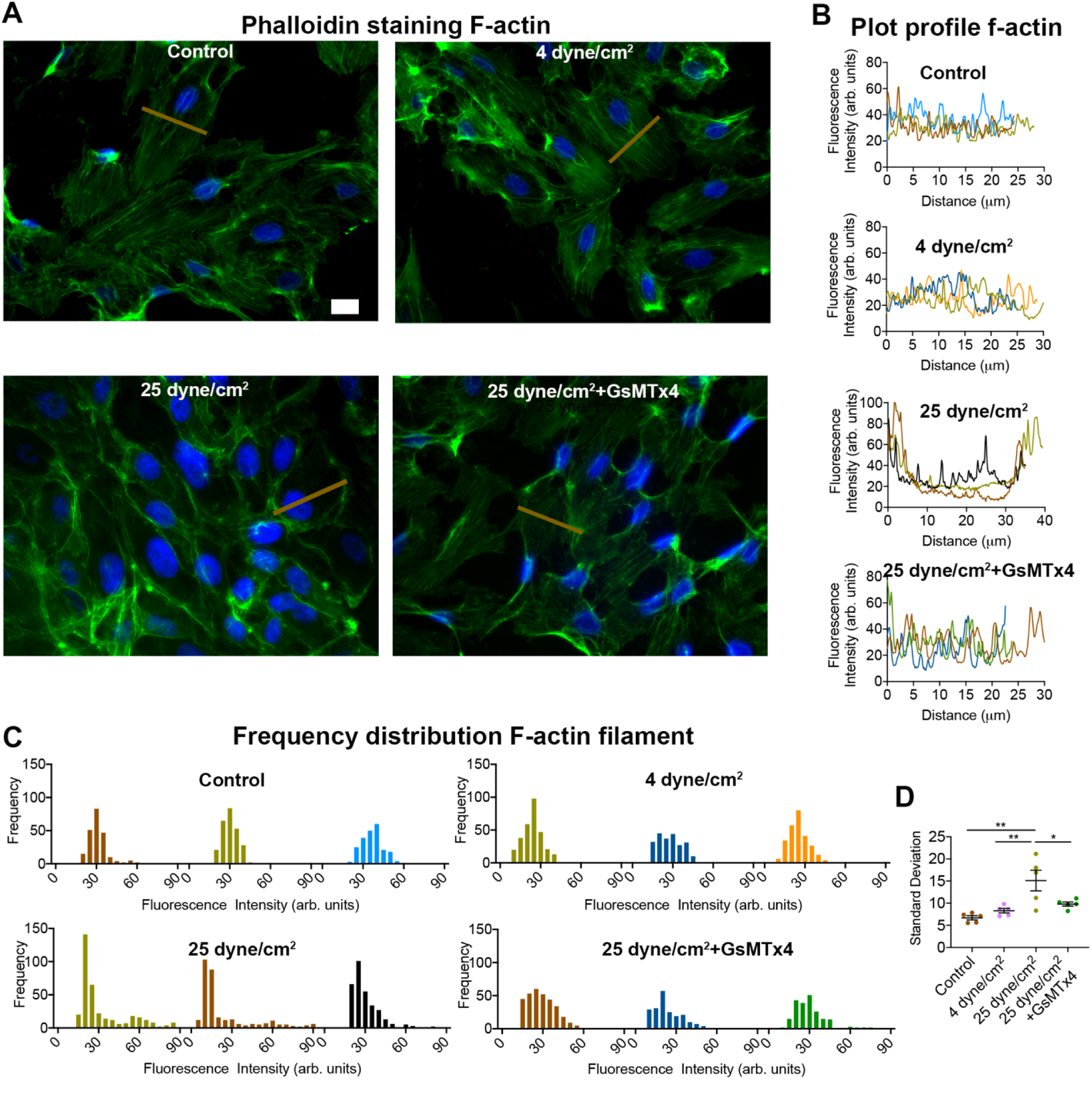
GsMTx4 inhibited F-actin disorganization in HUVECs caused by high shear stress. (A) Representative images show F-actin filament organization (green) and distribution in HUVECs under culture conditions of no shear stress (control panel), or shear stress applied for 10 min at 4 dyne/cm2, and 25 dyne/cm2 with or without GsMTx4(10 μM). Data are representative of three experiments. (B) Plot profile of F-actin from three cells. As represented in A, lines (tan color) were drawn perpendicular to F-actin orientation across which intensity profiles for Alexa 488 phalloidin were measured. (C) The distribution frequency of Alexa 488 phalloidin fluorescence intensity to measure F-actin distribution is shown for data in panel B. (D) The standard deviations of the grouped intensity frequency data measuring the dispersion from the mean intensity are shown from data in panel C. Individual data points shown represent mean ± SEM; Statistical analyses were performed using 1-way ANOVA with Tukey’s multiple comparisons. *P ≤ 0.05; ** P ≤ 0.01. Scale bar: 10 μm.

### Piezo1 mediated F-actin disruption depends on TRPV4 activation

To determine if activation of Piezo1 can affect the F-actin disorganization and might be the mechanism through which shear stress causes cytoskeletal remodeling, we applied the Piezo1 agonist, Yoda1 (10 μM) for 10 min to HUVECs. Without Yoda1 administration (control), the F-actin fibers are parallel in orientation and evenly distributed. Application of Yoda1 (10 μM) caused a drastic change in HUVEC morphology with apparent cell contraction and reduction in cell surface area, complete loss of F-actin fiber distribution, and accumulation of high-density clusters of F-actin fibers at the periphery of cells (Fig. 5). These Yoda1-induced effects were prevented by the TRPV4 blocker, HC067 (1μM), confirming that Piezo1 mediated F-actin disorganization required TRPV4 activation (Fig. 5). We determined that arachidonic acid (10 μM), an endogenous mediator of TRPV4 activation, triggered the same pattern of F-actin disorganization and gathering of high-density clusters of F-actin fibers in the periphery of HUVECs (Fig. 5). These findings signify that TRPV4 activation can cause cytoskeleton damage, but the shear stress-triggered remodeling of F-actin and disruption of F-actin fibers results from the combined actions of Piezo1 and TRPV4.

**Figure 5.**
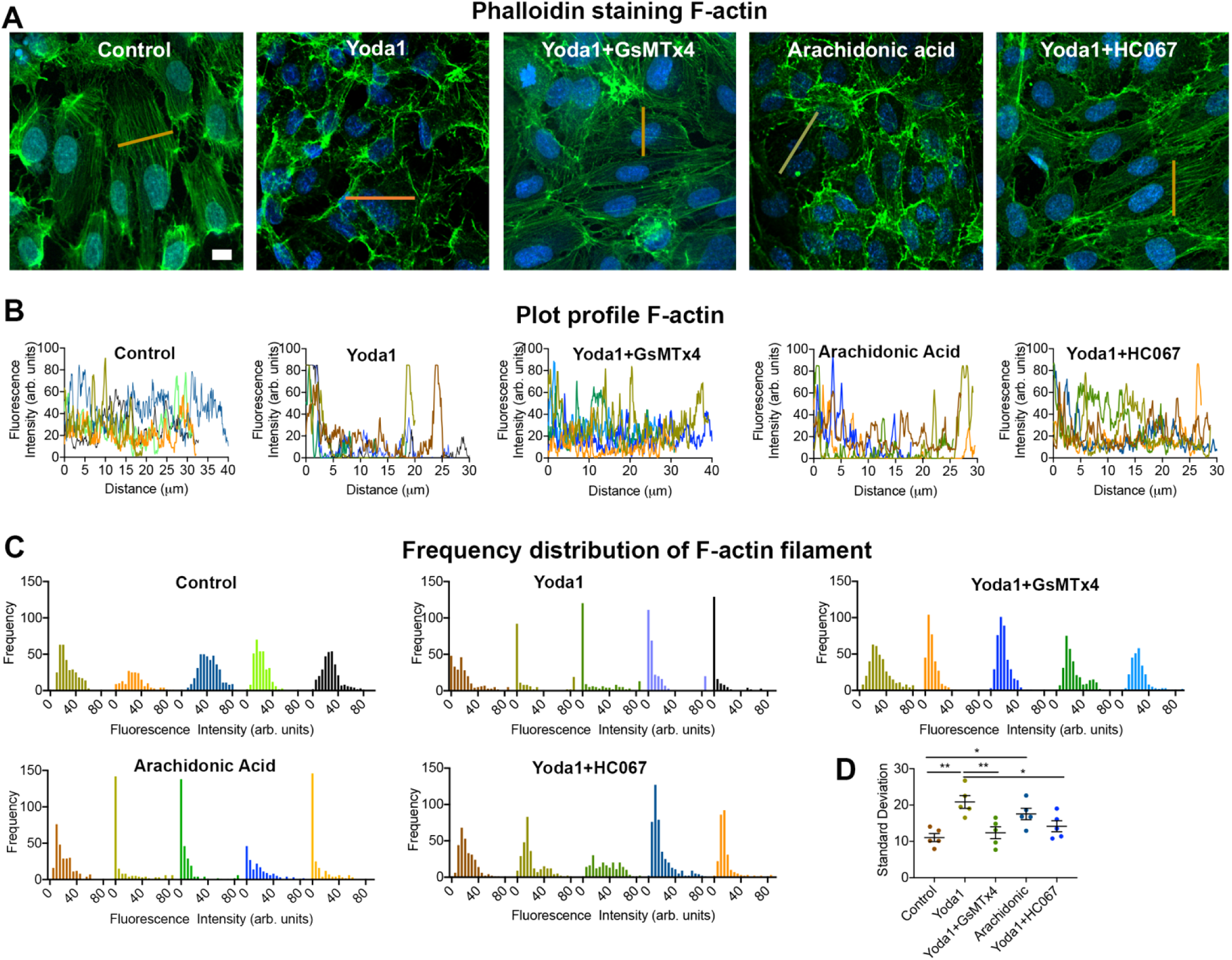
GsMTx4 and HC067 protect against Yoda1-induced F-actin disruption in HUVECs. Representative images of HUVECs show the distribution of F-actin filaments (Alexa 488 phalloidin, green) 10 min after treatment with DMSO (control), Yoda1 (10 μM), Yoda1 (10 μM) + GsMTx4 (10 μM) and Yoda1 (10 μM) + HC067 (5 μM). Images are representative of three experiments. (B) Plot profile of F-actin from five cells. A line was drawn perpendicular to filament orientation for plot profile analysis of F-actin and intensity profiles of Alexa 488 phalloidin across these lines were measured. (C) The fluorescence intensity frequency distribution of Alexa 488 phalloidin (to measure F-actin distribution) across the lines measured of data B. (D) The standard deviation of the grouped intensity frequency data measuring the dispersion from mean intensity of data C. Individual data points shown with mean ± SEM; Statistical analyses were performed using by 1-way ANOVA with Tukey’s multiple comparisons. *P ≤ 0.05; ** P ≤ 0.01. Scale bar: 10 μm.

### Blockade of TRPV4 inhibits Piezo1-mediated monocyte adhesion to endothelial monolayers

Disruption of normal cell-to-cell contacts in the vascular endothelium accompanies excessive shear force and leads to an inflammatory response characterized by monocyte adhesion (3,31). Having demonstrated that shear force activates Piezo1 which can disrupt AJs and form paracellular gaps in confluent monolayers we next evaluated the effects of Piezo1 on endothelial cell activation and adhesion protein expression, which are two of the early processes in atherosclerosis (41). Exposing endothelial cells to Yoda1 (1 μM) for 10 h increased expression of the adhesion protein VCAM1 and stimulated the attachment of monocyte cells (THP-1 cells) (Fig. 6). THP-1 attachment was prevented by the TRPV4 antagonist HC067 (Fig. 6, A and B). These findings indicate that the adverse effects of Piezo1-induced changes in endothelial cell adhesion lead to the attraction of inflammatory cells and are mediated by TRPV4.

**Figure 6.**
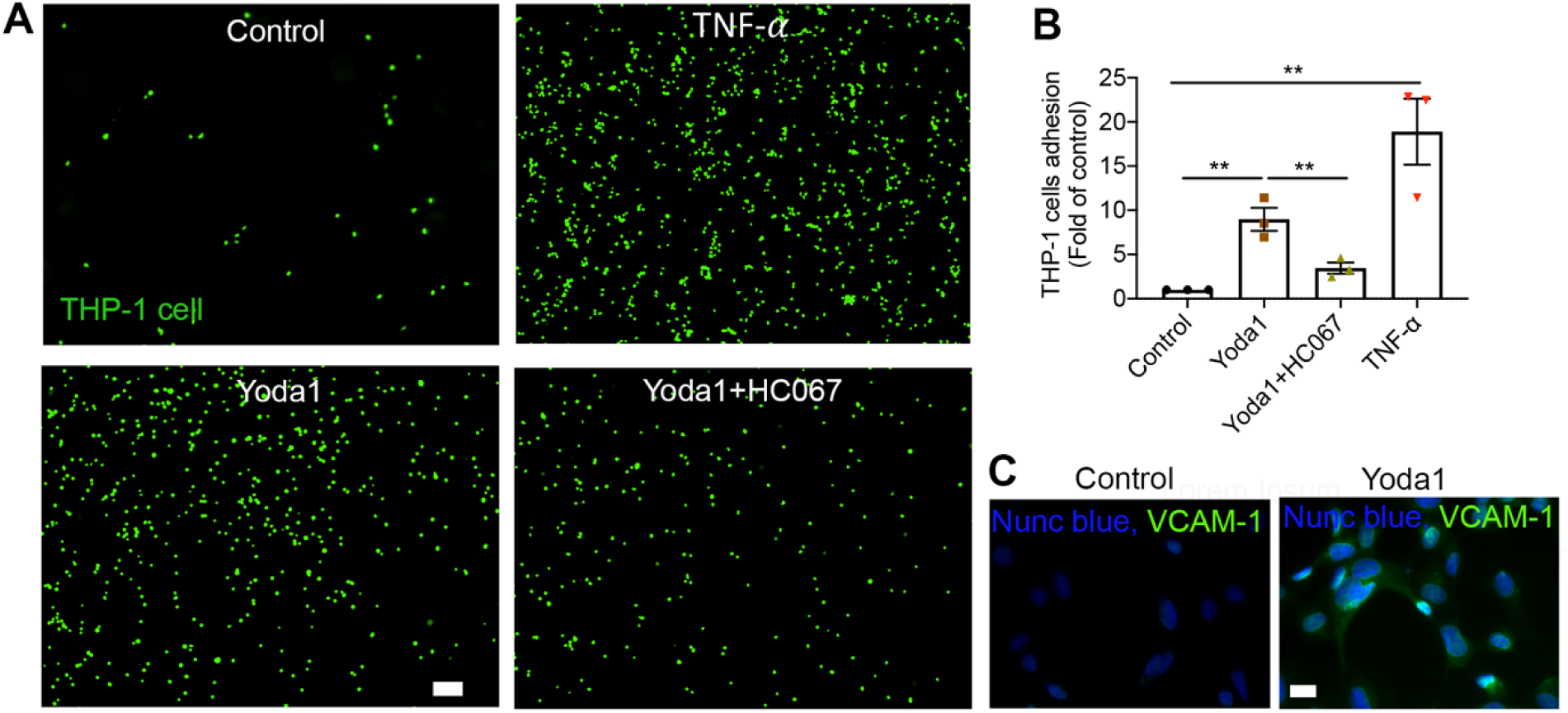
HC067 prevents the Yoda1 mediated THP-1 cell adhesion to HUVECs monolayer. (A) The representative images showing the HC067 (1 μM) prevent the Piezo1 agonist Yoda1 (1 μM) mediated THP-1 cells (green) adhesion to HUVECs monolayer. HUVECs monolayers were incubated with DMSO (control), Yoda1, Yoda1+HC067 and TNFα (10 ng/ ml) as a positive control for 10 h, and then THP-1 cells were added for 1 h to attach. Scale bar: 100 μm (B) The quantification of data A from three independent experiments. (C) The expression of VCAM1 (green) in endothelial cells with and without Yoda1 (1 μM) for 10 hr. Nuclei stained with Nunc Blue and Scale bar: 10 μm. Individual data points are shown with mean ± SEM; Statistical analyses were performed using by 1-way ANOVA with Tukey’s multiple comparisons. ** P ≤ 0.01.

## Discussion

Calcium signaling in endothelial cells is a highly regulated process (31). Under physiological conditions, the amplitude and frequency of [Ca^2+^]_i_ oscillations in endothelial cells maintain cellular integrity, but stimulation by inflammatory mediators such as histamine, bradykinin and thrombin increases intracellular calcium up to 10 fold (31). High sustained [Ca^2+^]_i_, leading to calcium overload, alters normal calcium signaling pathways and causes cytoskeletal disorganization and AJs disassembly. Together these processes facilitate transendothelial migration of leukocytes.

High blood pressure and shear stress also have been associated with endothelial dysfunction (1,3,13,31,42). High blood pressure or shear stress-mediated calcium overload contributes to cytoskeletal disorganization, AJs disassembly and leukocyte transendothelial migration (2,10,11,17,29,40). However, the severity of damage is endothelium-dependent, as the sensitivity to shear stress differs between the arterial and venous systems (43). Endothelial cells sense pressure and shear forces through calcium permeable mechanosensitive ion channels that convey extracelluar signals to maintain cellular integrity. The extent of calcium influx depends upon the strength of blood flow shear stress (44) and the number of mechanosensitive ion channels in the particular endothelium. We demonstrated in HUVECs that prolonged high shear stress produced a sustained elevation in [Ca^2+^]_i_ in contrast to low levels of shear stress or transient high shear stress. HUVECs are typical of other endothelial cells and express two types of mechanosensitive ion channels, Piezo1 and TRPV4, which have been implicated in vascular pathophysiological conditions (10,11,17,45,46). Either Piezo1 or TRPV4 may contribute to abnormally high [Ca^2+^]_i_ and may even initiate pathological events in endothelial cells (46-48). Piezo1 has a low single-channel conductance of approximately 22 pS and is a fast inactivating channel (15). Therefore, it seemed unlikely that Piezo1 alone could produce the prolonged elevation in [Ca^2+^]_i_ to induce a calcium overload state necessary to disrupt endothelial cell integrity. This led us to search for another calcium signaling pathway or calcium permeable ion channel that might be involved. TRPV4 was a likely candidate due to its slow inactivation kinetics and high single-channel conductance of approximately 60 pS (49). However, unlike Piezo1, TRPV4 does not appear to possess direct mechanosensing properties (50).

Although TRPV4 is sensitive to physical forces, such as shear stress, osmotic pressure, and mechanical stretching (4,22,25,51), the mechanical gating of TRPV4 has not been described. Though shear stress and cell swelling activate TRPV4 channel activity, TRPV4 senses mechanical force indirectly. During shear stress, TRPV4 is activated by 5,6 -EET, which is an endogenous ligand derived from arachidonic acid through the activation of PLA2 (26,28). Recently, we discovered that high pressure-induced Piezo1 activation caused the stimulation of PLA2 and subsequent opening of TRPV4 in pancreatic acinar cells (28). HUVECs sense mechanical force in a similar manner. In HUVECs, shear stress stimulates TRPV4 channel opening through Piezo1-stimulated activation of PLA2. Binding of calcium ion to PLA2 accelerates enzyme activity, which initiates the arachidonic pathway, 5,6 -EET production, and TRPV4 channel activation. It appears that a threshold level of peak [Ca^2+^]_i_ was necessary to induce PLA2 activation. It is likely that peak [Ca^2+^]_i_ is controlled by the number of Piezo1 channel openings. It could be that brief shear stress activated only a subset of Piezo1 channels resulting in submaximal [Ca^2+^]_i_ elevation which was insufficient to activate PLA2 in endothelial cells. In contrast, Yoda1 which would be expected to activate most Piezo1 channels, mimicked the effects of prolonged high shear stress-mediated [Ca^2+^]_i_ elevation.

Basal endothelial calcium levels control vascular permeability and contraction. Notably, basal calcium levels are altered in hypertension and high shear stress, which leads to disruption of the cell-cell junctional protein VE-cadherin and loss of endothelial barrier function. Impaired endothelial barrier function contributes to atherosclerosis which is a major cause of mortality among patients with hypertension. It was recently reported that blocking Piezo1 or TRPV4 prevented hypertension-associated vascular hyperpermeability but the underlying mechanisms or possible interactions were not uncovered (10,11). Nevertheless, our findings demonstrate how TRPV4 may contribute to micro-endothelial permeability (10,29). It has been shown that TRPV4 knockout mice are protected from lung vascular hyperpermeability in response to high pulmonary venous pressure (PVP) (20,51-53). Similarly, pharmacological blockade with the TRPV4 antagonist GSK2193874 also protected against pulmonary venous pressure (PVP)-induced permeability of the lung (10). It was recently reported that Piezo1 senses high vascular pressures at the lung endothelial surface and is responsible for vascular hyperpermeability and pulmonary edema (11). Thus, it appears that the high PVP-mediated vascular hyperpermeability is due to both Piezo1 and TRPV4. Our results demonstrated that Piezo1 activation caused the loss of VE-cadherin at the cell-cell junction and disrupted endothelial barrier integrity which were prevented by TRPV4 blockade. Moreover, TRPV4 was responsible for Piezo1 initiated cell retraction and paracellular gap formation in monolayer cultures. Thus, the combined actions of both Piezo1 and TRPV4 may contribute to the endothelial vascular barrier dysfunction in hypertension.

Fluid shear stress caused by blood flow is a major determinant of vascular remodeling and arterial tone and can lead to the development of atherosclerosis (31,40). High shear stress-mediated calcium signaling activates endothelial cells, cytoskeleton remodeling, and production of inflammatory cytokines and chemokines which facilitate leukocyte rolling, adhesion, and transendothelial migration (31). Our results suggest that all of these effects can be initiated by Piezo1 activation. Therefore, loss of endothelial barrier integrity and monocyte adhesion which are integral to atherosclerosis may be directly related to Piezo1 (19,48). Although our findings provide insight into the relationship between Piezo1 activation and deleterious effects on endothelial cells, it is important to note that development of atherosclerosis is a chronic process and is best studied in animals. Therefore, the relationship between Piezo1 and TRPV4 in the endothelium and initiation and perpetuation of atherosclerosis is more complicated than what can be studied *in vitro*. Nevertheless, the effects of Piezo1 on endothelial cell integrity may be an early step in the disease.

Piezo1 initiated the remodeling in our *in vitro* system. It is possible that a similar mechanism occurs under conditions of elevated shear stress *in vivo*. Due to the rate of blood flow, the endothelium in the arterial system is exposed to higher shear stress forces compared to the venous endothelium. In addition, transient or pulsatile shear stress is beneficial in the arterial system (2). In our experiments, using cells of venous origin, high shear stress of 25 dyne/cm2 for minutes at a time caused adverse effects. Regional distribution of both Piezo1 and TRPV4 throughout the vascular endothelium has not been extensively evaluated. It is possible that deleterious effects of high pressure do not occur in some vascular beds due to the absence of coexpression of Piezo1 and TRPV4 in the same cell. Our findings suggest that cells lacking either Piezo1 or TRPV4 would be protected from high shear stress-induced changes in cytoskeleton, cell traction, and inflammatory cell adhesion. Similarly, pharmacological blockade of either Piezo1 or TRPV4 would be expected to prevent the deleterious effects of shear stress in endothelium. However, identification and functional chacterization of both Piezo1 and TRPV4 throughout the vascular endothelium would be needed before considering therapeutics for cardiovascular diseases directed against these mechanosensitive ion channels.

The deleterious effects of shear stress in endothelial cells are secondary to sustained Ca2+ influx through TRPV4 channels. Therefore, the extent of Ca2+ influx could be influenced by several factors including the strength and duration of mechanical force, the level of PLA2 activity, and TRPV4 abundance. Our findings suggest that Piezo1 or TRPV4 alone is not sufficient to transduce mechanical force into pathological events. We have previously demonstrated that coupling of Piezo1 with TRPV4 is responsible for the effects of high pressure in pancreatic acinar cells (28). If functional coupling between Piezo1-TRPV4 is a generalized process in which TRPV4 translates the mechanical force sensed by Piezo1 into pathological events, then other organ systems like urinary bladder and bone tissue where Piezo1and TRPV4 are coexpressed (21,54-57), may also be amenable to targeting Piezo1 or TRPV4 to prevent adverse effects of pressure or shear stress.

## Experimental procedures

### Cell Culture

Primary human umbilical vein endothelial cells (HUVECs) (CC-2519) were obtained from Lonza. Cells were cultured with Endothelial Cell Growth Media (Cell Applications; 211-500) in a humidified incubator at 37°C and 5% CO_2_ (58). Cells were fed every two days and were not allowed to grow to confluence unless required for a specific experiment. Cells were used for experiments between passage numbers 2 to 5. HUVECs were passaged with 0.025% Trypsin/EDTA solution (TE, #R001100, Thermo Fischer Scientific). THP-1 cells, a monocyte cell line, were obtained from ATCC; TIB-202 (59). The suspension culture of THP-1 was grown in RPMI 1640 + sodium pyruvate (1 mM) + HEPES (10 mM) + glucose (10 mM) + 10% FBS + 0.05 mM beta-mercaptoethanol in a humidified incubator at 37°C and 5% CO_2_. Like HUVECs, the THP-1 cells were fed every two days and during culture, the cell number was not allowed to exceed 1 × 106. Primocin (InvivoGen) at a concentration of 100 μg/mL was added to cell culture media.

### Calcium Imaging

HUVECs were plated on a thin layered Matrigel coated plate for 1 h to allow cells to attach. The cells were then loaded with Calcium 6-QF (Molecular Devices) in a Minimum Essential Medium:Hanks’ balanced salt solution (MEM:HBSS; 1:1) for 30 min at 37°C in a CO_2_ incubator (27). The cells were washed gently with HBSS buffer with 2 mM Ca2+. The cells were imaged by using a Zeiss Axio observer Z1 with a 20x objective and images were captured at 400 ms intervals. HBSS buffer with 2 mM Ca2+ was used during imaging. The intracellular calcium elevation of individual cells was analyzed with MetaMorph software (Molecular Devices). Cells overloaded with dye or only faintly fluorescent were excluded from analysis. The chemicals used in calcium imaging experiments included: Yoda1 (Tocris; 5586), GsMTx4 (Abcam; ab141871), 5, 6-eicosatrienoic acid (Santa Cruz Biotechnology; sc-221066), arachidonic acid (Sigma; A3611), HC067047 (Tocris; 4100), GSK1016790A (Sigma; G0798), AACOCF3 (Tocris; 1462), YM26734 (Tocris; 2522).

### Monocyte-endothelial cell adhesion assay

To evaluate Yoda1-induced monocyte (THP-1 cells) adhesion to HUVECs, confluent HUVEC monolayers were pretreated with DMSO (vehicle control), Yoda1 (1 μM), Yoda1 (1 μM) + HC067 (1 μM), or TNF-α (10 ng/mL), (CELL BIOLAB. INC; Catalog; CBA210) for up to 10 hours in a humidified incubator at 37°C and 5% CO_2_. After treatment, the media were removed and THP-1 cells were added to HUVEC monolayers for 1 hour at 37°C to allow attachment (60). Before THP-1 cells were added to HUVEC monolayers, the THP-1 cells were harvested and prepared in a cell suspension at 1.0 × 106 cells/mL in serum free media and loaded with LeukoTracker (a live cell imaging dye) from CELL BIOLAB. INC; CBA210) as per manufacturer instructions. The non-attached THP-1 cells were cleared by gently washing three times with serum-free media. The remaining adherent THP-1 cells loaded with LeukoTracker dye were imaged using a Zeiss Axio observer Z1 with a 4x objective. Adherent cells were counted using FIJI software. The average number of THP-1 cells were calculated from five image frames. The data presented are representative of three independent experiments.

### Immunofluorescent Staining

HUVEC cells were washed with PBS (pH 7.4) and then fixed with 4% PFA for 10 min at room temperature. The fixed cells were treated with 0.1% Triton X-100 in PBS (pH 7.4) for 15 min at room temperature then blocked with 1% bovine serum albumin (BSA) for 1 h at room temperature. Cells were incubated with antibodies against VE-cadherin (Santa Cruz Biotechnology; sc9989) overnight at 2-8°C or antibodies against VCAM1 (abcam; ab134047) for 1 h at room temperature (61). Secondary goat anti-rabbit or secondary goat anti-mouse IgG Alexa Fluor 594 (Jackson ImmunoResearch) was used after each step for 1 h at room temperature. Immunostaining of F-actin was performed on fixed cells using Alexa Flour 488TM Phalloidin (Thermo Fisher scientific; A12379) for 1 h at room temperature. Cells were then washed three times with PBS for 10 min and incubated with Nunc Blue (Thermo Fisher scientific; R37606) and then mounted onto microscope slides with ProLong® Gold antifade reagent (Invitrogen). Images of immunostained cells were taken with a Zeiss Axio observer Z1 with a 40x objective or a Leica SP5 inverted confocal microscope with a 40x oil objective. The images were processed initially with MetaMorph software (Molecular Devices) or Leica LAS AF lite software and then the intensity of F-actin and width of VE-cadherin were measured with FIJI software (11).

### Shear stress experiments

Laminar flow shear stress was generated using parallel-plate fluid flow chambers from Ibidi, GmbH. Two types of flow chambers were used in the experiments. A μ-Slide I 0.4 Luer flow chamber was used to achieve 12 dyne/cm2 or lower shear stress. A μ-Slide I 0.2 Luer flow chamber was used for forces of 25 dyne/cm2 (17,28). The constant fluid flow rate with shear stress (τ) was determined as follows: τ = η × 104.7 f for μ-Slide I 0.4 Luer and τ = η × 330.4 f for μ-Slide I 0.2 Luer, where η = viscosity of the medium and φ = flow rate (according to the manufacturer’s instructions, Ibidi).

### Statistical analysis

Results are expressed as mean ± SEM. Mean differences between 2 groups were analyzed by 2-tailed Student’s *t*-test, and mean differences between multiple groups were analyzed by 1-way ANOVA with Tukey’s multiple comparison post-test (GraphPad Prism 8). *P* values of less than 0.05 were considered significant.

## Data availability

All the data described are contained within the manuscript and associated supporting information.

## Authors contributions

SMS and RAL designed experiments. SMS performed the experiments and analyzed data. SMS and RAL wrote the manuscript. RAL supervised the project and provided funding for the project.

## Acknowledgements

The authors thank Joelle Romac and Steven Vigna for helpful ideas during the course of this work and reviewing the manuscript. This work was supported by NIH grants R01 DK109368, R01 DK120555, and the Department of Veterans Affairs BX002230.

## The abbreviations used are

HUVEC: human umbilical vein endothelial cell
TRPV4: transient receptor potential vanilloid subfamily 4
PLA2: phospholipase A2
AJs: adherens junctions
5′,6′-EET: 5′,6′-epoxyeicosatrienoic acid
HC067: HC067047
GSK101: GSK1016790A
PVP: pulmonary venous pressure

